# Optogenetic activation of parabrachial tachykinin1 neurons drives nonphotic circadian entrainment

**DOI:** 10.64898/2026.08.17.745358

**Authors:** Victor Y. Zhang, Sekun Park, Kimberly D. Derderian, Jordan L. Pauli, Richard D. Palmiter, Horacio O. de la Iglesia

**Author notes:** Correspondence (Victor Y. Zhang), (Richard D. Palmiter), and (Horacio O. de la Iglesia). these authors contributed equally to this work (Victor Y. Zhang, Sekun Park).

## Abstract

Mammalian circadian rhythms are primarily entrained by light, but nonphotic cues can also reorganize behavioral timing through mechanisms that remain poorly understood. Nocturnal foot shocks delivered to rodents while they forage away from the safety of their nesting area have been shown to entrain circadian behavioral rhythms and shift foraging and feeding to the daytime. To identify the neural circuits underlying this nonphotic fear entrainment, we optogenetically stimulated tachykinin 1-expressing neurons in the parabrachial nucleus (Tac1^PBN^) during the subjective night while the animals foraged outside of their nest, which recapitulated the total activity-rest phase switch in circadian behaviors induced by foot shocks. Furthermore, selective stimulation of Tac1^PBN^ projections to the central amygdala (CeA) produced a significant but reduced phase shift compared to direct stimulation of Tac1^PBN^ cell bodies. When *Bmal1*, a core clock gene, was conditionally deleted from the CeA, mice failed to fear-entrain, implicating the CeA molecular clock as a necessary component for fear entrainment. Together, these experiments demonstrate that activation of a defined neuronal population outside of the suprachiasmatic nucleus (SCN) can reorganize circadian behavior by engaging a non-SCN circadian oscillator network that requires an intact CeA molecular clock.

## INTRODUCTION

Time is one of the most important dimensions of a species’ ecological niche, defined as the temporal coordination of organismal functions to optimize survival and reproduction. Shaped by prominent environmental cycles, endogenous biological rhythms represent a central feature of the temporal niche. In mammals, the expression of circadian rhythms is regulated by a distributed network of biological oscillators organized under the hierarchical control of a central pacemaker located in the suprachiasmatic nucleus (SCN). The daily light–dark (LD) cycle is the dominant environmental cue, or zeitgeber, that entrains the SCN and aligns circadian rhythms of behavior and physiology to the external world^1–3^. Nonetheless, nonphotic cues can also act as zeitgebers (e.g., cycles of food availability or ambient temperature)^4–9^, and these cyclic stimuli engage neural and molecular mechanisms that are distinct from those mediating photic entrainment of the SCN^6,10–13^. Compared to photic entrainment, the neural mechanisms supporting nonphotic entrainment remain poorly defined.

Field studies in free-living species have demonstrated that predator pressure can reorganize the 24-h daily activity pattern of animals. Specifically, prey species often reduce encounters with predators by shifting their activity away from periods of greatest risk, altering the temporal niche^14–19^. Despite the prevalence of these predator-induced temporal niche shifts in the wild, the mechanisms that translate recurring threat into stable changes in behavioral timing remain unknown. Because these shifts appear to represent behavioral plasticity within individuals^14^, rather than through intergenerational changes (e.g., natural selection), they raise the possibility that repeated threat engages the circadian system itself to predict safer times for foraging and other behaviors.

Recent work has shown that cyclic fear acts as a salient nonphotic zeitgeber that entrains circadian rhythms of behavior^20,21^. These findings were obtained using custom-designed cages that simulate a more naturalistic environment, in which freely moving mice move between a safe nesting compartment and a foraging area with *ad libitum* food and water access where foot shocks can be delivered through a metal grid floor. When foot shocks are randomly distributed during the light phase (light fear, LF), animals exhibit enhanced nocturnal foraging and feeding relative to baseline. In contrast, when foot shocks occur during the dark phase (dark fear, DF), animals shift their circadian foraging and feeding activity into the daytime. Remarkably, this diurnal activity pattern persists under free-running conditions (constant darkness without shocks), demonstrating the existence of one or more fear-entrainable oscillators (FrEOs) that encode the temporal structure of nocturnal fear. Importantly, although the SCN molecular clock is necessary for fear entrainment, it is not sufficient and remains stably entrained to the light–dark (LD) cycle^14^, indicating that non-SCN FrEOs that communicate with the SCN can override the SCN’s photic entrainment to drive behavioral rhythms.

A neural population well positioned to convey the timestamp of recurring threat to the circadian system would both carry the threat signal itself and project to structures capable of translating that signal into a change in circadian timing. The parabrachial nucleus (PBN) is a general brainstem alarm hub that relays diverse threat signals ^22^, including foot shock, to the forebrain principally through the spino-parabrachio-amygdaloid pathway^23–25^. Genetically distinct PBN populations are recruited by foot shock, with activation of Tac1-^26^ and CGRP-expressing^27,28^ neurons reproducing distinct components of defensive behaviors spanning active (escape) and passive (freezing) coping. These populations project densely to the central amygdala (CeA) and bed nucleus of the stria terminalis (BNST)^24,29^, limbic targets that mediate affective behavior and display SCN-dependent circadian clock gene rhythms ^30–32^, making the PBN a plausible entry point for interrogating the circuitry underlying fear entrainment.

Here, we show that time-specific activation of Tac1^PBN^ neurons is sufficient to drive a total switch in circadian activity-rest phase that persists under free-running conditions, whereas activation of CGRP^PBN^ neurons drives aversion without entrainment. Stimulation of Tac1^PBN^ terminals in CeA, but not the BNST, partially recapitulated this effect. Conditional deletion of the clock gene *Bmal1* in CeA abolished fear entrainment, implicating a local CeA molecular clock as a necessary component of the fear-entrainable oscillator network.

## RESULTS

### Tac1^PBN^ activation is sufficient to entrain circadian foraging rhythms

To test whether local activation of PBN alarm circuitry is sufficient to entrain circadian behavior, we developed an optogenetic fear-entrainment chamber modeled on the foot shock-based paradigm. The chamber continuously recorded nesting, foraging, and feeding activity with infrared sensors while animals were housed under constant darkness (DD). A microswitch at the transition between the nesting and foraging compartments gated optogenetic stimulation only when the animal entered the foraging area (Fig. 1). Thus, optogenetic stimulation was delivered with the same temporal and spatial contingency as nocturnal foot shocks in a previous study^20^, only during a defined 12-h subjective-night window and only when mice occupied the aversive foraging compartment.

**Figure 1.**
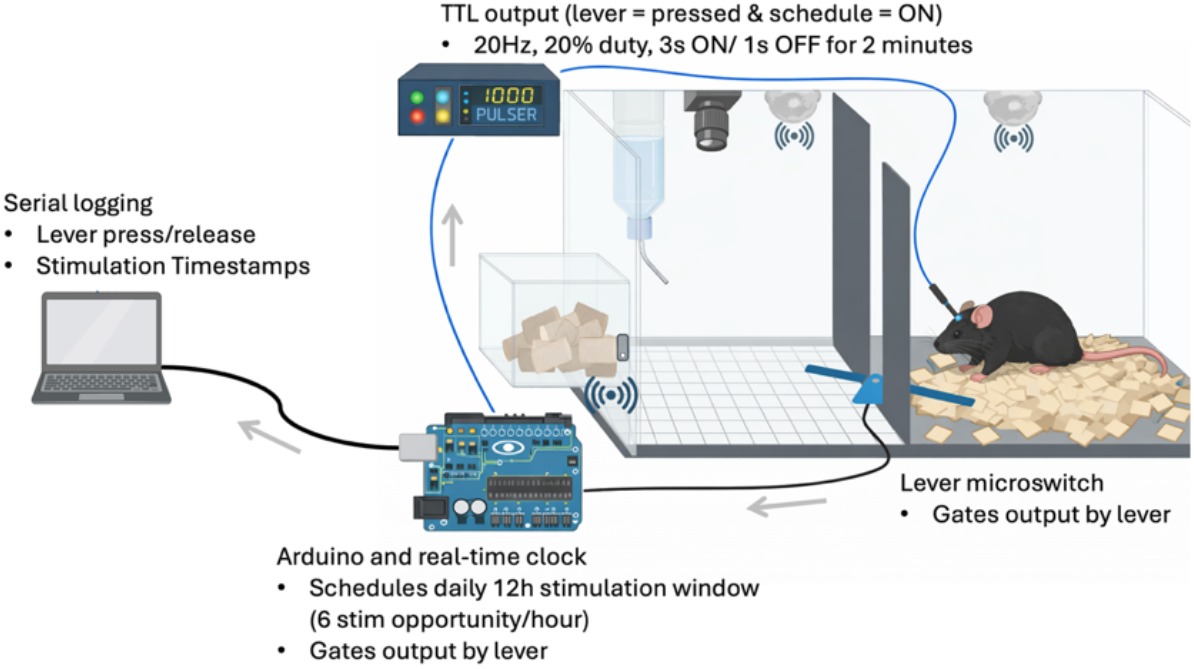
A schematic of the optogenetic fear-entrainment chamber. Mice, tethered to fiber optics, were allowed to move freely between a nesting (cage right) and a foraging area (cage left), which contained *ad libitum* food and water, via a corridor composed of a balanced pivot lever connected to a microswitch. The microswitch provided lever press information to an Arduino, which gates closed-loop optostimulation so that laser pulses are only delivered while an animal occupies the foraging area. Infrared beams record nesting, foraging, and feeding activity.

Aversive stimuli recruit *Tac1* neurons in the external lateral PBN (elPBN), many of which overlap with calcitonin gene-related peptide (CGRP)-expressing neurons^26^. Despite this anatomical overlap, Tac1^PBN^ and CGRP^PBN^ neurons appear to support distinct defensive strategies, with activation of Tac1^PBN^ neurons being sufficient for driving active coping, whereas activation of CGRP^PBN^ neurons drives passive coping^26^. Therefore, we wondered which of these behavioral responses (or both) is required for fear entrainment. *Calca*^Cre/+^ (encodes CGRP) or *Tac1*^Cre/+^ mice received bilateral elPBN injections of AAV_DJ_-DIO-ChR2-YFP and optic fibers over the PBN (Fig. 2a–c). Control mice received AAV_DJ_-DIO-YFP, with fibers placed over either PBN or CeA, and were pooled for subsequent analyses. Slice histology confirmed viral transduction and fiber placement in ChR2-expressing PBN neurons (Fig. 2a-c).

**Figure 2.**
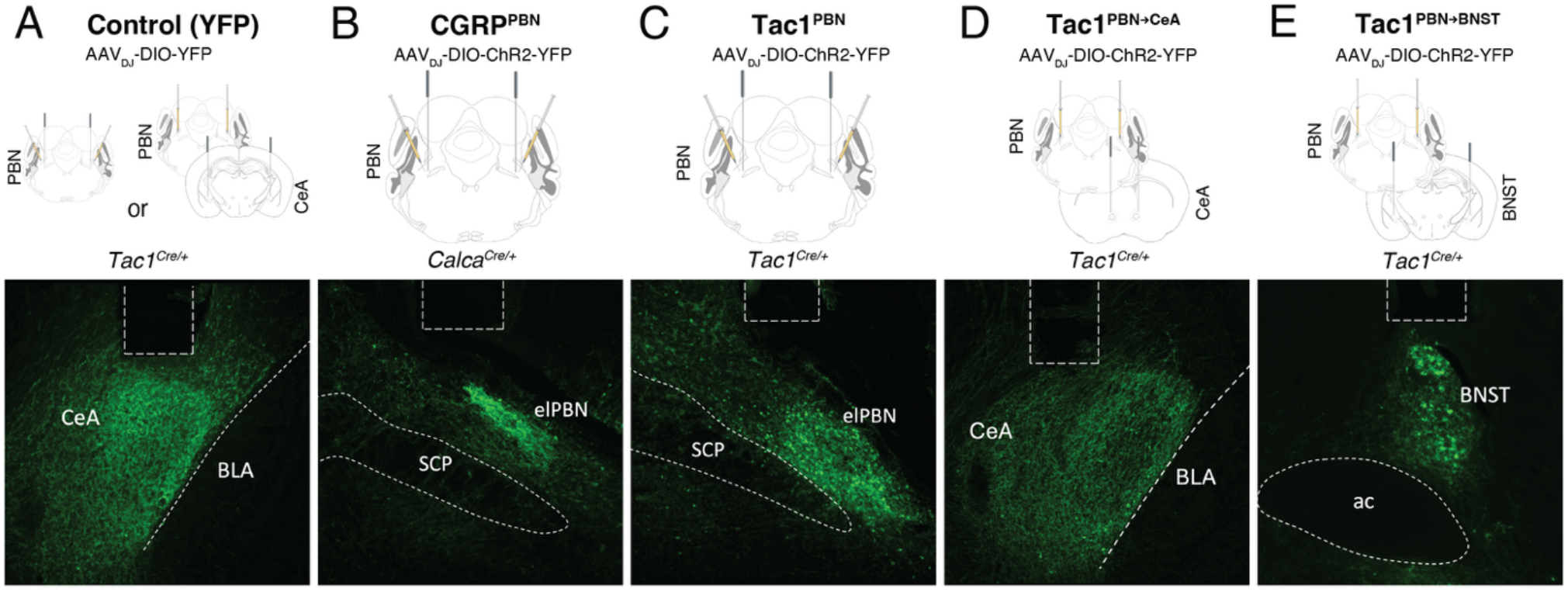
Viral targeting and optic fiber placement for optogenetic stimulation of PBN populations and their axon terminals. For all panels, top: schematic of viral injection and fiber placement; bottom: representative coronal section immunolabeled for YFP or ChR2–YFP (green), with the fiber tract outlined by dashed lines. (A) Cre-dependent YFP (AAV_DJ_-Ef1α-DIO-YFP) was expressed bilaterally in the PBN of *Tac1*^*Cre/+*^ mice, with optic fibers positioned over either the external lateral PBN (elPBN) or the CeA; groups were pooled for subsequent analyses (Control, n = 5). The representative image shows YFP-labeled Tac1^PBN^ terminals in CeA beneath the fiber tract. (B, C) Cre-dependent ChR2 (AAV_DJ_ -Ef1α-DIO-ChR2-YFP) was expressed bilaterally in PBN of *Calca*^*Cre/+*^ (B; CGRP^PBN^, n = 6) or *Tac1*^*Cre/+*^ (C; Tac1^PBN^, n = 6) mice, with optic fibers over the elPBN. Representative images show ChR2–YFP-expressing cell bodies in the elPBN. (D, E) Cre-dependent ChR2 was expressed PBN of *Tac1*^*Cre/+*^ mice, with optic fibers positioned over the CeA (D; Tac1^PBN→CeA^, n = 6) or the BNST (E; Tac1^PBN→BNST^, n = 5). Representative images show ChR2–YFP-labeled Tac1^PBN^ axon terminals in CeA (D) and BNST (E) beneath the fiber tract. ac, anterior commissure; BLA, basolateral amygdala; SCP, superior cerebellar peduncle.

After recovery, mice were transferred to optogenetic fear-entrainment chambers in DD. Following 10 d of baseline recording, mice were exposed for 10-14 d to a 12-h window of optostimulation opportunity during the subjective night, from circadian time 12 (CT12; activity onset) through CT24. Following this stimulation phase, mice were released into constant free-running conditions with no optostimulation for 10 days. Optostimulation trains (20 Hz, 15-ms pulses, 3 s ON/1 s OFF, 2-min maximum duration) were scheduled once every 10 min at random times during the subjective night but could only be delivered while the animal was in the foraging area. If the animal retreated to the nest, ongoing stimulation terminated immediately (Fig 1).

Optostimulation induced escape from the foraging area in all ChR2 groups compared to YFP controls (Cox proportional hazards model, Tac1^PBN^: HR = 79.1, P < 0.001; CGRP^PBN^: HR = 23.0, P < 0.001; Fig. 3). The response was fastest in Tac1^PBN^ mice, which retreated with a median latency of 3 s after effective stimulation onset, whereas CGRP^PBN^ mice showed a slower response with a median latency of 10 s (Fig. 3). Thus, stimulation of either PBN population was acutely aversive, but Tac1^PBN^ activation produced the most rapid termination of foraging. Moreover, the two PBN populations had distinct effects on daily activity patterns (Fig. 4). During the stimulation phase, Tac1^PBN^ mice redistributed foraging activity away from the subjective night and into the subjective day (Fig. 4c). This shift was evident in animal waveforms (Fig. 4d-f) of foraging activity and diurnality – defined as activity during the baseline subjective day– measures, with Tac1^PBN^ mice approaching near-complete subjective daytime foraging during the stimulation phase (diurnality = 98.7 ± 2.5%; Fig. 5a). In contrast, CGRP^PBN^ stimulation reduced the amplitude of subjective-night foraging (Fig. 4e) but did not produce a comparable nocturnal-diurnal switch (diurnality = 43.2 ± 4.5%; Fig. 5a).

**Figure 3.**
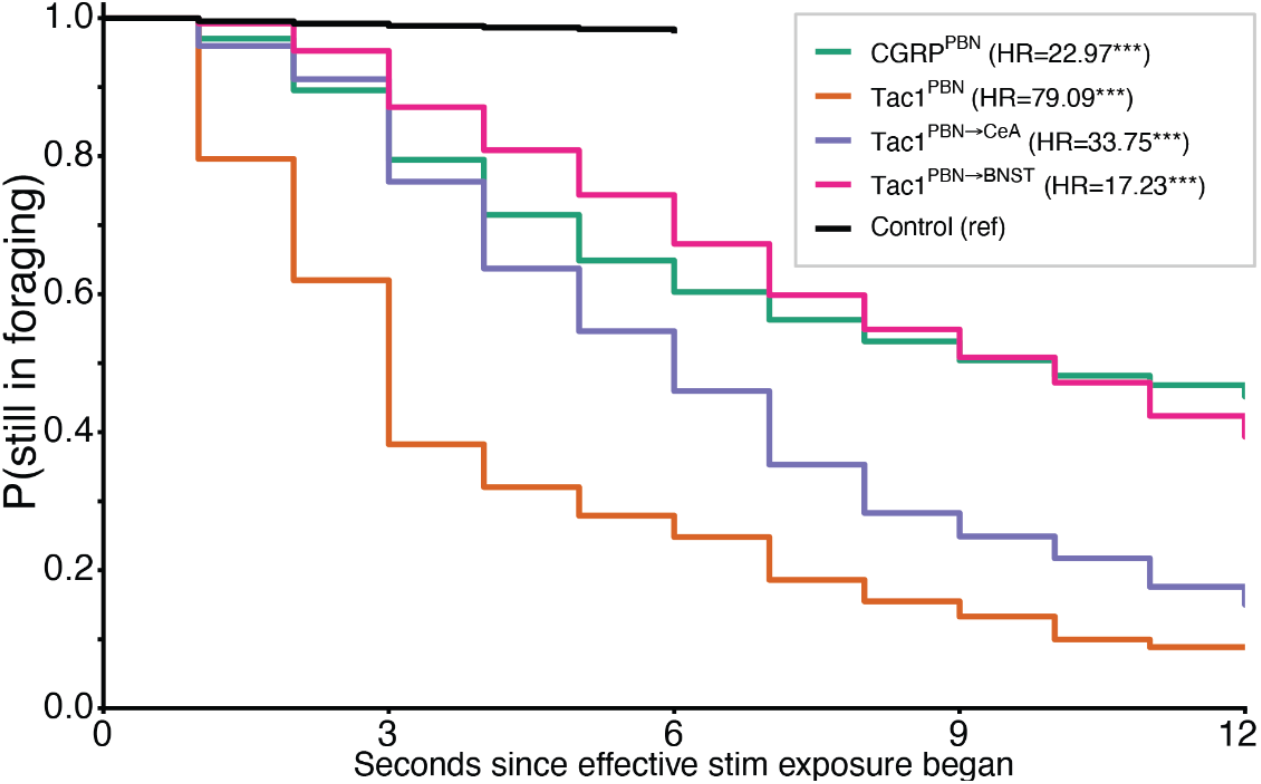
Acute escape from the foraging area during optostimulation. Kaplan–Meier probability of remaining in the foraging area after effective optostimulation onset. Hazard ratios of opsin-expressing mice versus YFP controls (Cox proportional-hazards model) are indicated in the legend. Control episodes were capped at a 6-s effective epoch and ChR2-expressing groups were followed to a 2-min maximum. Group comparisons were performed in a matched 6-s window. ***P < 0.001. n = 5 YFP control, 6 CGRP^PBN^, 6 Tac1^PBN^, 6 Tac1^PBN→CeA^ and 5 Tac1^PBN→BNST^ mice.

**Figure 4.**
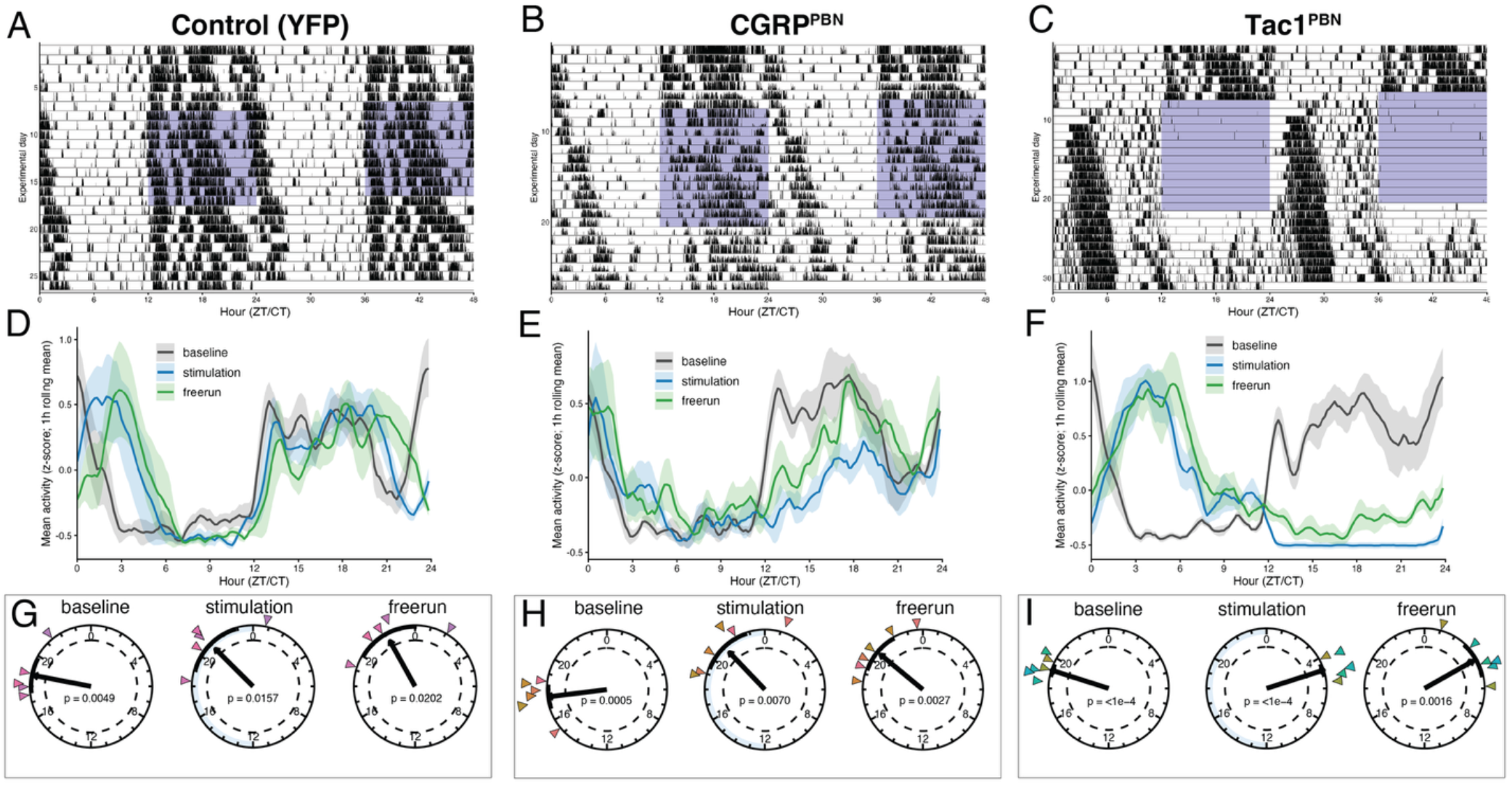
Time-specific optogenetic activation of Tac1^PBN^, but not CGRP^PBN^ neurons drives entrainment of circadian behaviors. (A–C) Representative double-plotted actograms of infrared foraging activity across baseline, optostimulation and free-run for a YFP control (A), a CGRP^PBN^ (B) and a Tac1^PBN^ (C) mouse; shaded regions mark the boundaries of the subjective-night (CT12–24) optostimulation window over 14 days. (D–F) Group-mean foraging waveforms (z-scored activity, 1-h rolling mean; mean ± SEM) for baseline, stimulation and free-run for YFP control (D), CGRP^PBN^ (E) and Tac1^PBN^ (F) mice. (G–I) Rayleigh plots of the daily center-of-gravity (COG) phase for each stage for YFP control (G), CGRP^PBN^ (H) and Tac1^PBN^ (I) mice; each colored triangle is a unique mouse, the bold vector is the group-mean resultant and the dashed circle marks the critical resultant length for P = 0.05, with 95% confidence intervals shown as black bands along the inner circumference. n = 5 YFP control, 6 CGRP^PBN^, 6 Tac1^PBN^ mice. n = 5 YFP control, 6 CGRP^PBN^, 6 Tac1^PBN^ mice.

**Figure 5.**
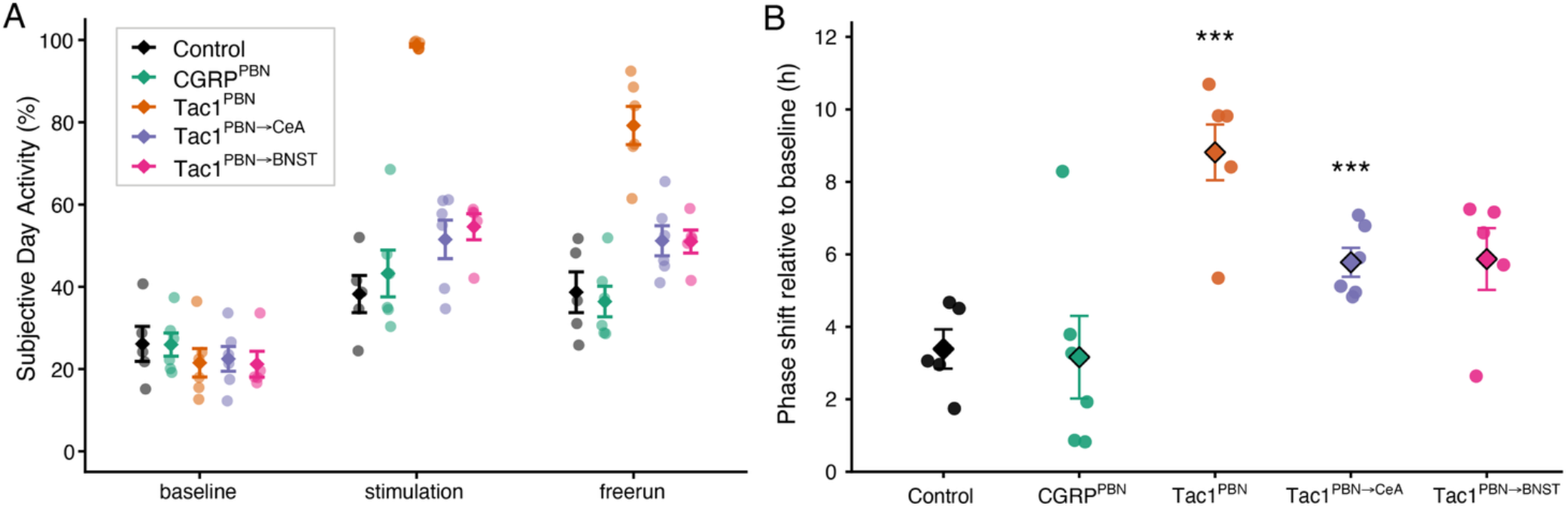
Subjective daytime activity and circadian quantification across optogenetic stimulation groups. (A) Subjective-day activity (diurnality index, %) across baseline, stimulation, and free-run. (B) Baseline-normalized, free-running phase shift. Group mean ± SEM is shown overlayed with individual mice. Asterisks denote pairwise circular permutation tests versus control (***P < 0.001, Holm adjusted); the shift relative to control mice was significant for Tac1^PBN^ and Tac1^PBN→CeA^ but not for Tac1^PBN→BNST^ or CGRP^PBN^. n = 5 YFP control, 6 CGRP^PBN^ 6 Tac1^PBN^, 6 Tac1^PBN→CeA^ and 5 Tac1^PBN→BNST^ mice.

The redistribution of activity during stimulation could reflect masking rather than a shift of the underlying pacemaker. In contrast to masking, true entrainment of a circadian oscillator requires that the activity shift remains after removal of the cyclic stimulus; thus, we examined behavior after optostimulation was discontinued. During this free-running phase, Tac1^PBN^ mice continued to show elevated subjective-day foraging, and the phase of foraging activity remained similar to that observed during stimulation (Fig. 4c, f). By contrast, CGRP^PBN^ mice recovered subjective-night activity during the free-running phase, consistent with an acute masking response rather than stable entrainment (Fig. 4b, e). Animals maintained body mass between the baseline stage and optostimulation stage across all experiments.

We quantified circadian phase using the daily center of gravity (COG) of foraging activity and analyzed phase shifts across stages with circular statistics. During baseline, YFP controls, Tac1^PBN^ mice, and CGRP^PBN^ mice all showed significantly clustered COG phases that reflected the previous LD cycle that animals were exposed to (controls: CT18.7±1.15 [mean±95% CI]; r = 0.94; Tac1^PBN^: CT19.2±0.43; r = 0.99; CGRP^PBN^: CT17.5±0.80; r = 0.96; Fig. 4g-i). During stimulation and free-running, COG phase was delayed from baseline in all groups, including controls (stimulation: controls CT21.0±1.92 (r = 0.86), Tac1^PBN^ CT4.8±0.43 (r = 0.99), CGRP^PBN^ CT21.1±1.74 (r = 0.85); free running: controls CT22.1±2.04 (r = 0.84), Tac1^PBN^ CT4.0±1.23 (r = 0.92), CGRP^PBN^ CT20.6±1.37 (r = 0.90); Fig. 4g-i). The modest delay in YFP controls likely reflects the intrinsic free-running period of the animals in DD and/or an effect of the blue laser light that turns on during the subjective night. To account for this shift over the course of each experiment, we compared baseline-normalized, free-running phase shifts between each ChR2 group and the YFP control group using two-sample permutation-based circular tests. Tac1^PBN^ stimulation produced a significantly larger free-running phase delay than controls (mean shift = 8.8 ± 0.8 h (mean ± SEM), Holm-adjusted P < 0.001; Fig. 5b). In contrast, the CGRP^PBN^ shift did not differ significantly from controls (mean shift = 3.2 ± 1.1 h, Holm-adjusted P = 0.76; Fig. 5b). Together, these data show that repeated, time-specific activation of Tac1^PBN^ neurons is sufficient to produce a stable, activity-rest phase switch in circadian foraging behavior, whereas activation of CGRP^PBN^ neurons produces acute aversion without circadian entrainment. Given this, we focused subsequent optogenetic experiments on Tac1^PBN^ neurons a model for nonphotic entrainment.

### Tac1^PBN^ projections to the limbic forebrain drive partial circadian phase shifts

We next asked whether nonphotic entrainment is mediated by downstream limbic targets of Tac1^PBN^ axons. Tac1^PBN^ neurons project prominently to the CeA and BNST^26,29^, both of which display circadian clock gene rhythms that depend on SCN input^30–32^. We bilaterally injected Tac1^Cre/+^ mice with AAV_DJ_-DIO-ChR2-YFP and implanted optic fibers over either CeA (Tac1^PBN→CeA^, n = 6) or BNST (Tac1^PBN→BNST^, n = 5) (Fig. 2d,e). Mice were then tested with the same baseline, stimulation, and free-running protocol described above.

Optostimulation of Tac1^PBN^ terminals in either CeA or BNST increased retreat from the foraging area relative to YFP controls (Tac1^PBN→CeA^: HR = 33.8, P < 0.001; Tac1^PBN→BNST^: HR = 17.2, P < 0.001; Fig. 3). Terminal-evoked escape was slower than that observed after Tac1^PBN^ soma stimulation and was similar to the response observed in CGRP^PBN^ mice, with median retreat latencies of 6 s for Tac1^PBN→CeA^ and 10 s for Tac1^PBN→BNST^ (Fig. 3). Thus, activation of either terminal field was sufficient to induce escape responses, but response time was slower than in Tac1^PBN^ soma activation.

During the stimulation phase, both terminal-stimulation groups reduced foraging during the subjective night and increased foraging during the subjective day (Fig. 6a-d). This redistribution resulted in changes in diurnality (Tac1^PBN→CeA^: baseline 22.5 ± 3.9% vs. stimulation 51.5 ± 3.9%, F_(1,12)_ = 32.6, P < 0.001; Tac1^PBN→BNST^: baseline 21.2 ± 3.2% vs. stimulation 54.6 ± 3.2%, F_(1,5)_ = 97.8, P < 0.001; Fig. 5a). Lever-defined foraging bouts showed a similar pattern, where Tac1^PBN→CeA^ mice made more nighttime than daytime forays at baseline, but this day-night difference was reduced or abolished during stimulation, primarily because nighttime foray rate decreased from 258.6 ± 39.8 to 83.8 ± 8.8 bouts (Supp. Fig. 1). Tac1^PBN→BNST^ mice showed similar changes in bout structure, with nighttime foray rate decreasing from 167.3 ± 27.3 to 68.5 ± 9.7 bouts (Supp. Fig. 1).

**Figure 6.**
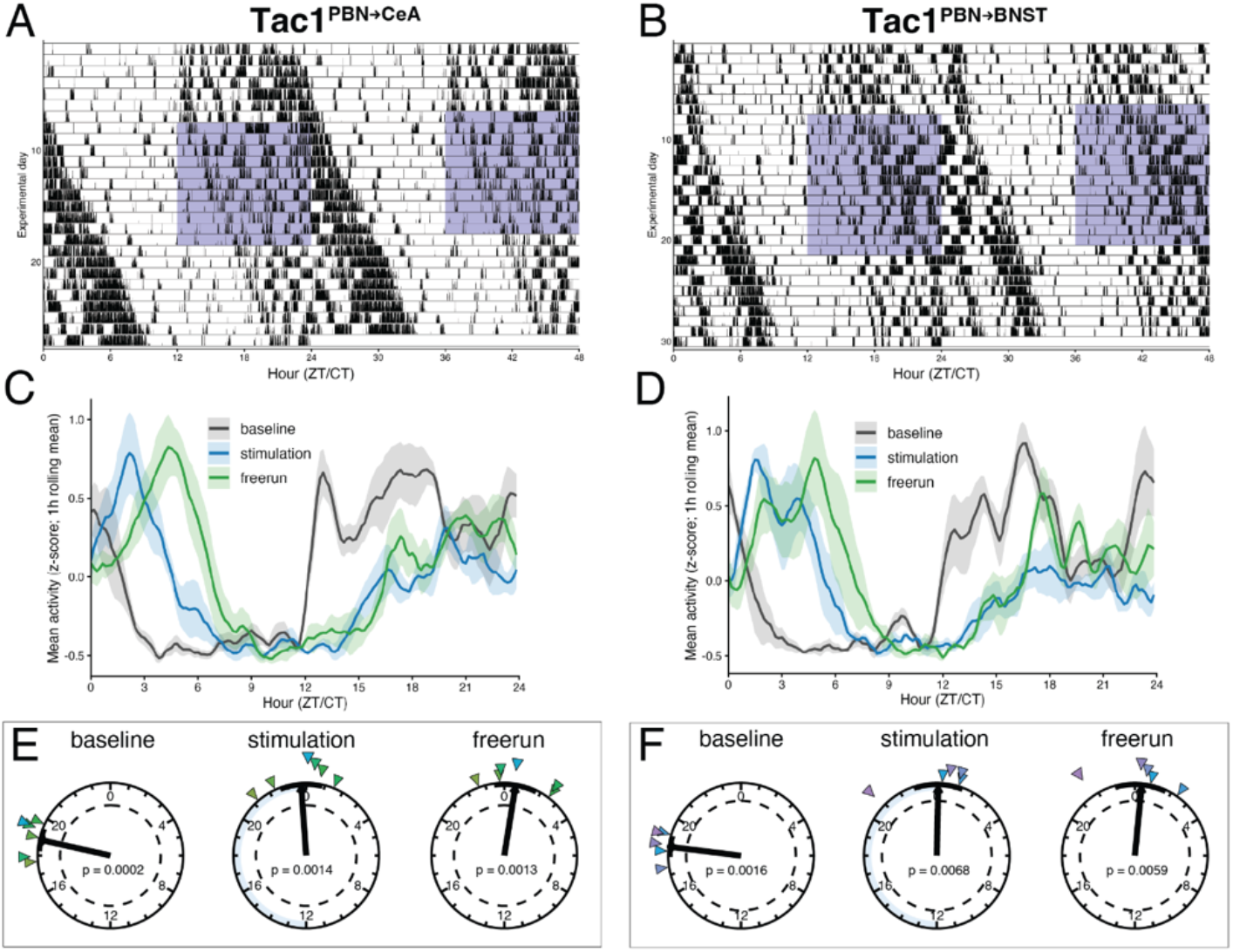
Stimulation of Tac1^PBN^ axon terminals in the central amygdala (CeA) and bed nucleus of the stria terminalis (BNST) produces partial circadian phase shifts. (A, B) Representative double-plotted actograms of infrared foraging activity across baseline, optostimulation and free-run for a Tac1^PBN→CeA^ (A) and a Tac1^PBN→BNST^ (B) mouse; shaded regions mark the boundaries of the nightly (CT12–24) optostimulation window. (C, D) Group-mean foraging waveforms (z-scored activity, 1-h rolling mean; mean ± SEM) for Tac1^PBN→CeA^ (C) and Tac1^PBN→BNST^ (D) mice. (E, F) Rayleigh plots of the daily center-of-gravity (COG) phase for each stage for Tac1^PBN→CeA^ (E) and Tac1^PBN→BNST^ (F) mice; each colored triangle is a unique mouse, the bold vector is the group-mean resultant and the dashed circle marks the critical resultant length for P = 0.05, with 95% confidence intervals shown as black bands along the inner circumference. n = 6 Tac1^PBN→CeA^ and 5 Tac1^PBN→BNST^ mice.

Post-stimulation, when animals were transferred to free-running conditions, terminal-stimulated mice maintained their phase-shifted patterns of foraging activity during the free-running phase (Fig. 6a, b). In both Tac1^PBN→CeA^ and Tac1^PBN→BNST^ groups, activity during the early subjective night remained reduced while activity during the early subjective day remained elevated (Fig. 6c, d), indicating that any underlying phase shift was likely driven by circadian entrainment rather than masking effects. COG-based circular analyses confirmed significant phase clustering across experimental stages for both terminal-stimulation groups (Rayleigh tests, all P < 0.05; Fig. 6e, f). Mean COG phase shifted from CT18.8±0.62 (r = 0.98) and CT18.5±0.54 (r = 0.99) at baseline to CT23.7±1.15 (r = 0.93) and CT0.1±1.33 (r = 0.92) during stimulation, and to CT0.6±1.16 (r = 0.93) and CT0.4±1.33 (r = 0.92) during free running, for Tac1^PBN→CeA^ and Tac1^PBN→BNST^ mice, respectively (Fig. 6e, f). Baseline-normalized, free-running phase shifts were larger than those of YFP controls but reached statistical significance only for the CeA projection (Tac1^PBN→CeA^: mean shift = 5.8 ± 0.4 h, Holm-adjusted P < 0.001; Tac1^PBN→BNST^: mean shift = 5.9 ± 0.9 h, Holm-adjusted P = 0.081; Fig. 5b).

Overall, these findings indicate that stimulation of Tac1^PBN^ terminals over CeA can shift circadian foraging behaviors, whereas stimulation of terminals over BNST was less effective. However, neither terminal field fully recapitulated the large active-rest phase inversions observed in direct Tac1^PBN^ soma activation (Fig. 5b). The smaller terminal-stimulation effects suggest that the full Tac1^PBN^ entrainment phenotype may require coordinated recruitment of multiple downstream targets, local PBN circuit interactions, or both.

### Bmal1 deletion in the central amygdala abolishes foot shock entrainment

Although stimulation of Tac1^PBN^ neurons recapitulates the circadian entrainment produced by cyclic fear, this finding does not establish the PBN as the FrEO. Instead, the PBN could function primarily as an input-processing node that conveys aversive timing information to the oscillator. Because previous studies have shown that cyclic aversive stimuli can entrain clock-gene expression in the CeA^32^, we hypothesized that the PBN relays the temporal pattern of aversive signals to the CeA, thereby entraining a CeA FrEO. To test this hypothesis, we conditionally deleted the core clock gene *Bmal1* in CeA and tested entrainment to cyclic foot shocks randomly presented during the nocturnal –subjective night– phase (0.2 mA). To accomplish this, mice with floxed *Bmal1* (*Bmal1*^fl/fl^) received bilateral CeA injection of a Cre-expressing virus (AAV1-CMV-Cre-GFP) to delete *Bmal1* locally (*Bmal1*-cKO^CeA^; n = 7), with C57BL/6J mice serving as wild-type controls (WT; n = 5) (Fig. 7a, b). Both groups underwent the same experimental protocol previously described in Bussi et al.^20^ (i.e., baseline, subjective night foot shocks, and free-running phase).

**Figure 7.**
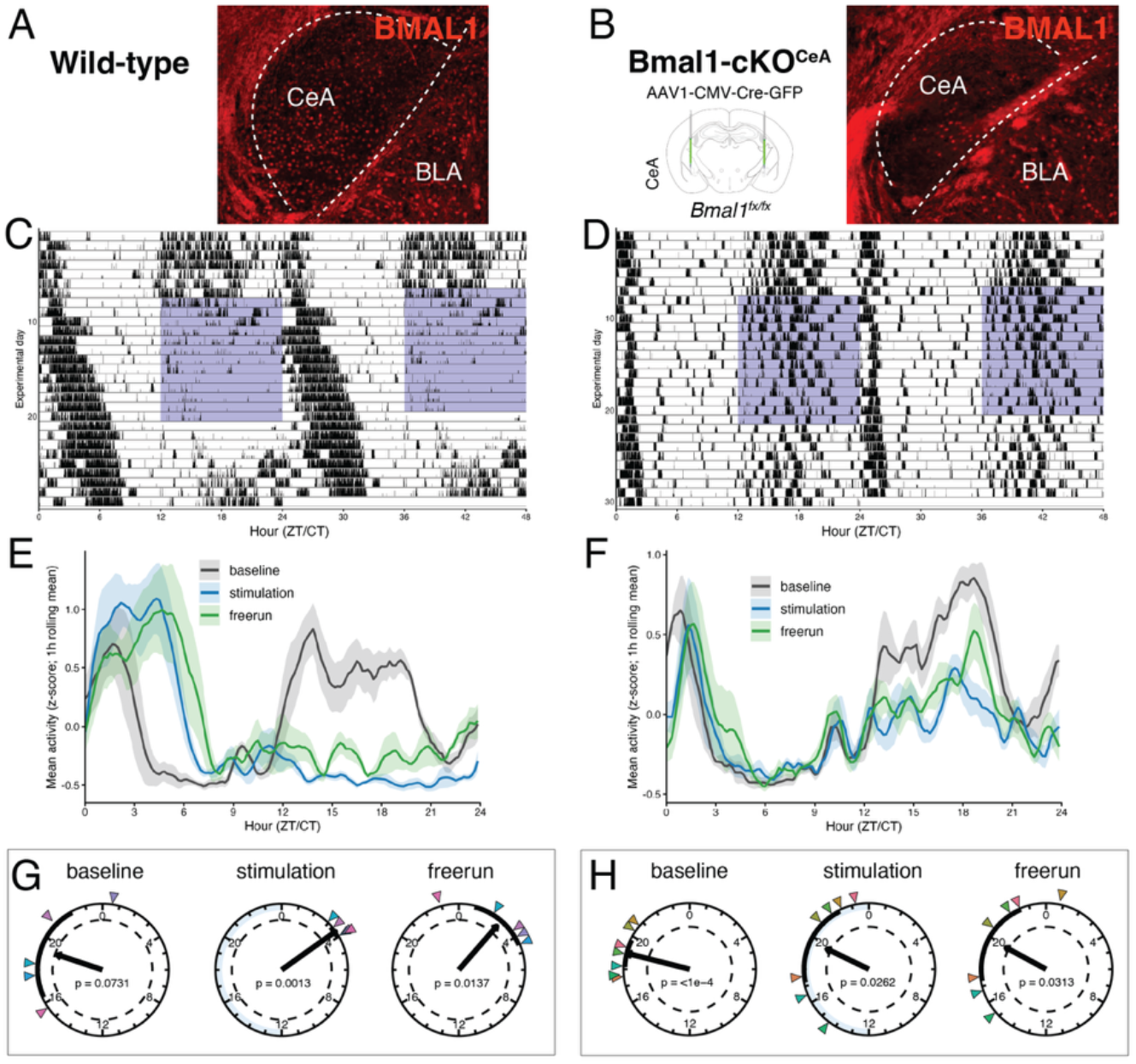
Conditional deletion of *Bmal1* in the central amygdala (CeA) abolishes foot-shock entrainment. (A, B) Representative coronal sections immunolabeled for BMAL1 (red) in a wild-type (A) and in a *Bmal1*^*fl/fl*^ mouse that received bilateral injections of AAV1-CMV-Cre-GFP into CeA to delete *Bmal1* in the CeA (*Bmal1*-cKO^CeA^), showing loss of BMAL1 immunoreactivity restricted to the CeA. (C, D) Representative double-plotted actograms for a wild-type foot shock control (C) and a *Bmal1*-cKO^CeA^ (D) mouse across baseline, foot shock and free-run; shaded regions mark the boundaries of the nightly (CT12–24) foot shock window (0.2 mA). (E, F) Group-mean foraging waveforms (z-scored activity, 1-h rolling mean; mean ± SEM) for WT (E) and *Bmal1*-cKO^CeA^ (F) mice. (G, H) Rayleigh plots of the daily center-of-gravity (COG) phase for each stage for wild-type (G) and *Bmal1*-cKO^CeA^ (H) mice; each colored triangle is a unique mouse, the bold vector is the group-mean resultant and the dashed circle marks the critical resultant length for P = 0.05, with 95% confidence intervals shown as black bands along the inner circumference. n = 5 WT and 7 *Bmal1*-cKO^CeA^ mice.

In WT controls, nocturnal foot shocks entrained circadian foraging behaviors to the subjective day. The COG phase of foraging shifted from CT19.2±2.91 (r = 0.71) at baseline to CT3.6±0.38 (r = 0.99) during the shock phase and remained day-shifted at CT2.7±1.77 (r = 0.87) under free-running conditions, with significant phase clustering during both the shock and free-running stages (Rayleigh P = 0.0013 and P = 0.014, respectively; baseline clustering was not significant).This was accompanied by an increase in subjective daytime activity from 34.9% during baseline to 89.7% during shocks and 74.4% during free-run (free-run vs. baseline within-group contrast, F_(1,5)_ = 38.2, P = 0.0016). In contrast, *Bmal1*-cKO^CeA^ mice failed to entrain to nocturnal foot shocks, remaining active during the subjective night throughout experimental stages (baseline CT19.0±0.83 [r = 0.96], shocks CT19.7±2.51 [r = 0.70], free-run CT19.9±2.59 [r = 0.68]); diurnality also remained low, moving only from 28.9% at baseline to 40.7% during shocks and 39.9% in free-run, never approaching the daytime redistribution seen in WT mice. Although the change in subjective daytime activity from baseline to free-running conditions reached significance within the *Bmal1*-cKO^CeA^ group (F_(1,7)_ = 14.5, P = 0.0066), this small change was likely due to free-running over a 14-d period and activity patterns still failed to invert to the subjective day (Fig. 7c-h and Fig. 8a).

**Figure 8.**
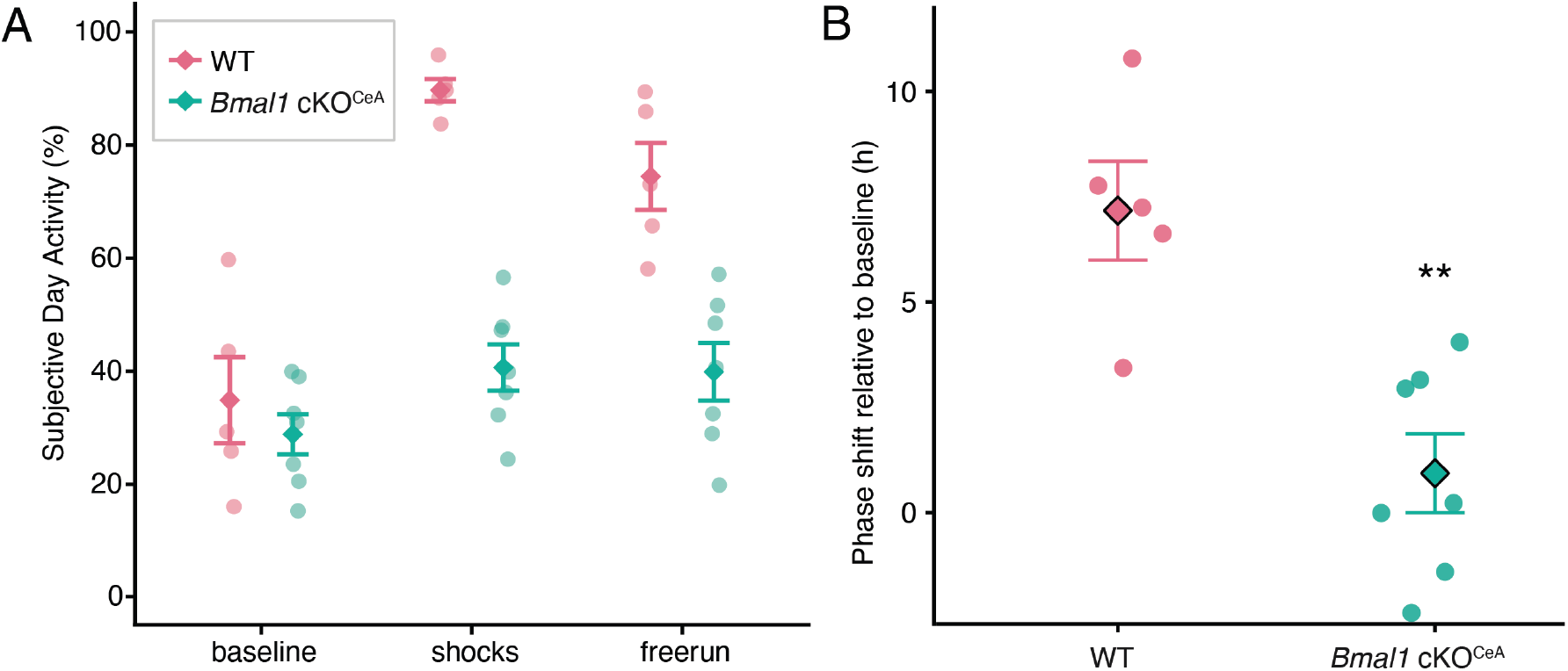
Mice with *Bmal1* deleted from the central amygdala (*Bmal1*-cKO^CeA^) exhibit less subjective daytime activity and shift their phase less than wild-type (WT) controls under conditions of fear entrainment. (A) Subjective-day activity (diurnality index, %) across baseline, foot shock, and free-run for WT and *Bmal1*-cKO^CeA^ mice. (B) Baseline-normalized free-running phase shift. Group mean ± SEM is shown overlayed with individual mice. (**P = 0.0032, circular permutation test, WT versus *Bmal1*-cKO^CeA^). n = 5 WT and 7 *Bmal1*-cKO^CeA^ mice.

Directly comparing the free-run and baseline activity phase difference between groups confirmed that the clock deletion abolished entrainment. The free-running phase shift was markedly smaller in *Bmal1*-cKO^CeA^ than in WT mice (0.9 ± 0.9 h vs. 7.2 ± 1.2 h; signed difference −6.25 h; P = 0.0032) (Fig. 8b). Together, these data indicate that an intact CeA molecular clock is required for foot shock entrainment, implicating CeA as a necessary fear-entrainable oscillator.

## DISCUSSION

The capacity of animals not only to respond to, but to anticipate, the predictable cycles (or “zeitgebers”) of their environment is the central adaptive function of circadian entrainment. Under predation pressure, prey species are known to routinely restructure their temporal niche, shifting the timing of their daily activity/rest cycles to avoid predators that hunt on a recurring schedule. Yet, how the mammalian circadian system encodes the timing of danger into an endogenously generated temporal program remains unknown. Here, we show that the PBN→CeA pathway serves as a key component involved in integrating and encoding the temporal structure of recurring threat to reshape circadian timing. We found that subjective night optogenetic activation of Tac1^PBN^ neurons was sufficient to drive a stable inversion of circadian foraging to the subjective day, whereas conditional deletion of the core clock gene *Bmal1* in the central amygdala (CeA), a major projection target of Tac1^PBN^ neurons, abolished foot shock entrainment, establishing Tac1^PBN^ activation as sufficient to drive entrainment and identifying the CeA molecular clock as necessary for it. These experiments demonstrate that targeted activation of a non-SCN population is sufficient to drive total inversions of activity-rest phase via nonphotic circadian entrainment.

Interestingly, our results indicate that acute aversion and circadian entrainment are dissociable. Activation of either Tac1^PBN^ or CGRP^PBN^ neurons elicited escape from the foraging area, yet only Tac1^PBN^ activation produced a persistent phase shift; CGRP^PBN^ activation suppressed subjective-night foraging during stimulation but did not redistribute activity into the day and recovered to baseline once stimulation ceased, consistent with an acute masking effect rather than entrainment. Eliciting a defensive response at a fixed circadian time is therefore not, by itself, sufficient to reorganize the circadian timing of behavior.

Previous studies have shown that a significant portion of CGRP^PBN^ neurons co-express Tac1^29^, and intersectional targeting of Tac1^+^;Calca^−^ neurons indicates that they can suppress some functions attributed to CGRP^PBN^ neurons, including conditioned taste version and freezing^26^. Nonetheless, the divergent responses between Tac1^PBN^ and CGRP^PBN^ activation suggest that the capacity to entrain circadian behavior is not a general consequence of PBN-mediated aversion or the expression of defensive behaviors.

We further demonstrate that nocturnal activation of Tac1^PBN^ axon terminals at individual downstream targets is sufficient to generate circadian phase shifts sufficient for generating partial subjective daytime activity. Efferents of the PBN are anatomically organized to distribute threat cues broadly across the extended amygdala^29,33,34^, structures whose integrity is required for fear entrainment^21^. Consistent with this organization, stimulating Tac1^PBN^ terminals in CeA produced significant partial phase shifts, whereas stimulating them in BNST produced a shift of comparable magnitude that did not reach statistical significance. However, neither reproduced the full inversion driven by activation of neuronal somas. Bowen et al.^24^ showed that CGRP^PBN^ neurons send collaterals to multiple brain regions and that activating individual terminals never recapitulated the effect of activating the cell bodies. Likewise, we suggest that somatic activation of Tac1^PBN^ neurons may recruit multiple targets in a coordinated manner that single-site terminal stimulation does not recapitulate.

Notably, the PBN, CeA, and/or BNST, could putatively function as a member of the FrEO network rather than as a non-clock relay to the FrEO. The CeA and BNST are especially strong candidates, as both sustain SCN-coupled clock-gene rhythms^30,31,35^, and an extra-SCN amygdalar oscillator has recently been shown to coordinate behavioral rhythms directly^36^. Because a node can be required to express an entrained rhythm without containing the oscillator that generates it, we tested this directly by conditionally deleting the molecular clock gene *Bmal1* in CeA, which abolished foot shock entrainment and supports CeA as a bona fide FrEO node rather than a non-clock relay. Future research should investigate the extent to which the PBN molecular clock is necessary for fear entrainment.

Previous work has shown that the SCN is necessary but not sufficient for fear entrainment, and its molecular clock remains locked to the light-dark cycle even as animals shift to daytime activity under cyclic nocturnal fear^20^, suggesting that the FrEO network ultimately overrides the light-entrained behavioral program. We propose two non-mutually exclusive hypotheses for this effect. First, a parabrachial-amygdala circuit could alter the physiological state of the SCN without shifting the molecular clock, as acute stress has been shown to shift the phase of SCN calcium activity^37^, and the SCN has also been shown to receive nonphotic afferents from limbic and autonomic structures, including BNST^38^. In addition, other limbic regions, such as the paraventricular thalamus (PVT), which is reciprocally connected with the SCN^39,40^ as well as other fear centers^41,42^, may act as conduits to relay circadian timing to the SCN^37,40^. Second, given the strong evidence that nocturnal-diurnal phase preference is determined downstream of the SCN clock^43–45^, FrEOs may instead be acting on SCN-coupled output circuits, with the rhythmic output of putative FrEOs competing with SCN-driven signals at shared downstream effectors of arousal/rest behavioral states (e.g., subparaventricular zone, dorsomedial hypothalamus, paraventricular nucleus of the hypothalamus, or ventrolateral preoptic area, among others).

Overall, our finding that individual axonal stimulation of downstream PBN targets produces only partial phase shifts supports a model of multiple, partially coupled and broadly distributed oscillators rather than a single central clock, paralleling other nonphotic circadian systems such as the food-entrainable system^12,46–48^. That said, the robust phase shifts produced by Tac1^PBN^ activation indicates that the FrEO network is tightly integrated within regions receiving input from the PBN. Compared to other modalities of nonphotic entrainment, such as entrainment by cyclic feeding, the FrEO network may be more easily tractable and consolidated within the limbic forebrain. Together, these findings highlight a neural mechanism enabling the reorganization of circadian behaviors in response to threatening stimuli, which is not only relevant to modeling predator-prey interactions but may also be relevant patients suffering from anxiety disorders and post-traumatic stress disorder for which circadian disruptions represent a core feature^49–52^.

## METHODS

### Animal subjects

Either *Tac1*^*Cre/+*^ (Jax # 021877) or *Calca*^*Cre/+*^ (JAX # 033168) mice were used for optogenetic stimulation experiments. For conditional deletion of *Bmal1* in the CeA, *Bmal1*^*fl/fl*^ (floxed-*Bmal1*) (Jax # 007668) mice and C57BL/6J wild-type controls were used. These experiments included animals of both sexes that were 8-12 weeks old at experimental onset. Mice were group housed with *ad libitum* access to food and water on a 12 h light/dark cycle at 21°C before experiments. After surgery, mice were singly housed. Viral expression was confirmed histologically at experimental endpoint, and only mice with correct targeting were included in the final analysis. All experimental procedures followed protocols approved by the Institutional Animal Care and Use Committee at University of Washington

### Surgical procedures

Mice were anesthetized with 5% isoflurane and head-fixed on a robotic (Neurostar) or manual stereotaxic frame, and 1.5-2% isoflurane was used to maintain anesthesia. Viral vectors were bilaterally injected using a 33-gauge blunt needle (NanoFil) or glass pipette (Nanoject II; Drummond Scientific) to the PBN (AP -4.90 mm, ML ±1.35 mm, and DV +3.40 mm) or CeA (AP -1.20 mm, ML ±2.90 mm, and DV +4.80 mm) at a rate of 0.1 µl/min until 0.3 µl had been infused. When optogenetic manipulation was planned, a fiber-optic cannula (RWD Life Science) was positioned just above the PBN (AP -4.90 mm, ML ±1.65 mm, DV 3.20 mm), CeA (AP -1.20 mm, ML ±3.00 mm, DV -4.50 mm), or BNST (BNST, AP +0.35 mm, ML ±1.15 mm, and DV -3.3 mm), and anchored with superbond (Loctite), C&B Metabond (Parkell) and dental acrylic (TEETS).

For optogenetic stimulation of PBN and CeA *Tac1* neurons, AAV_DJ_-Ef1α-DIO-ChR2-YFP virus was used for experimental group, and AAV-DIO-YFP virus for the control group were used. For conditional deletion of *Bmal1* in the CeA, AAV-CMV-Cre-GFP (Addgene 105545) was used. Virus titers were adjusted to approximately 10^12^ gc/ml. A recovery period of 4 weeks preceded any behavioral testing.

### Behavioral paradigm/chambers

#### Optogenetic chambers

Animals were housed in standard (19 × 40 × 18 cm [W × L × D]) mouse cage unless otherwise indicated. The cages (26 x 47.6 x 15.2 cm) used in optogenetic experiments were designed to enable automated, conditional stimulation of freely moving individually housed mice via tethered fiberoptics (Fig. 1). The nesting (containing corncob bedding) and foraging areas (containing ad libitum access to food and water) were equally divided by a black acrylic wall with a corridor (4.75 × 24 cm [W × L]) that provided access between the two areas. The filter top of each optogenetic chamber had a 1.5-cm slit cut through the midline that extended across nesting and foraging areas to allow for tethered animals to move freely between sides without obstruction. The corridor was designed as a balanced pivot lever connected to a microswitch that provided input to an Arduino, allowing detection of animal transitions between foraging and nesting areas. The Arduino provided input to a local computer (for serial logging of microswitch state) and a laser pulser (for delivering a stimulation program gated by the microswitch). Two passive infrared IR detectors recorded the activity within the nesting and foraging areas, respectively, and a break-beam IR sensor recorded nose-pokes into the food container, allowing us to continuously record the nesting, foraging, and feeding activity for each animal. All sensor data were acquired and stored in 1-min bins to a local computer using ClockLab software and hardware (Actimetrics, Wilmette, IL).

#### Foot shock chambers

Foot shock experiments were carried out in smaller chambers that shared the same layout as in the optogenetic experiments (as described in Bussi et al.^20^). Briefly, cages measuring 26 cm wide x 47.6 cm long x 15.2 cm deep were divided into two equal parts by a black acrylic wall to make two independent fear chambers. Each independent fear chamber, which was further divided into equally sized nesting and foraging areas, housed a single animal. The floor of the foraging area consisted of a metal grid connected to a precision shocker (Coulbourn Instruments, Allentown, PA) that was controlled by an Arduino programmed to activate the shocker within a prespecified temporal window (see *Experimental design*). Cages were fit with IR detectors as described above to record nesting, foraging, and feeding activity.

### Experimental design

Before each experiment, animals were entrained to a 12:12 LD cycle. Animals were then transferred to fear chambers. The fear entrainment protocol consisted of three different phases, all occurring in constant darkness (DD). First, a baseline phase (10-14 days) during which animals were allowed to become accustomed to the new cage chamber and during which baseline activity was collected from each animal. Animals were then subjected to the shocks or optostimulation phase, which lasted 10-14 days. In the *Bmal1*-cKO^CeA^ vs. WT experiments, the foraging area was rendered dangerous for a 12-h interval via random foot shocks during the subjective night (active phase of animal). The foot shocks consisted of activating the shock grid at 0.2 mA for 10 s randomly within every 20-min interval. In the optogenetic experiments, the foraging area was similarly rendered dangerous for 12 h; however, rather than foot shocks, animals were exposed to random blue light stimulations during the subjective night (the animal’s active phase). The optogenetic epochs (20 Hz, 15-ms pulse width, 3s ON/1s OFF), which were programmed to last 2 min, consisted of activating the laser pulser conditionally when the animal was in the foraging area (microswitch pressed) randomly within every 10-min interval. The stimulations were gated by the microswitch so that if the animal retreated to the nesting area at any point (microswitch released), the optogenetic stimulations would immediately terminate. Following the shock or stimulation phase, animals were released into constant conditions with no further shocks or stimulations for 10-14 days to determine the free-running phase of their circadian rhythms. Following the free-running phase, animals were transitioned to a regular 12:12 LD cycle and perfused at CT15.5.

### Immunohistochemistry and image acquisition

For histological verification, mice were deeply anesthetized with 5% isoflurane and perfused intracardially with cold PBS, followed by cold 4% PFA, and decapitated. To preserve fiber tracts, the skull around the cerebellum and olfactory bulbs was removed, and the entire head was placed in a 24 h post fix in 4% PFA at 4 °C. The next day, the brains were removed and further fixed in 4% PFA for 2-4 h and then placed in 30% sucrose for at least 3 d at 4 °C to allow for complete infiltration. Brains were then placed in a mold with OCT and frozen at −80 °C until sectioning. Sections were collected on a cryostat (Leica) at 35 µm for the regions of interest (PBN, BNST, CeA) and placed into a sucrose-based cryoprotectant for long term storage at −20 °C.

To stain the sections in order to amplify the signal from viral injections and better visualize projections, sections were first washed twice in PBS to remove the cryoprotectant before being moved into a blocking solution at room temperature containing 0.2% Triton-X in PBS with 3% normal donkey serum for 1 h. Sections were then moved into primary antibody in blocking solution (chicken-anti-GFP 1:10,000, Abcam, ab13970 and rabbit-anti-cFos 1:500, Abcam, ab) overnight at 4 °C. The next day, sections were rinsed in PBS 3 times for 10 min each and then placed in secondary antibody in PBS (Alexa Fluor 488 donkey anti-chicken 1:500, Jackson ImmunoResearch, AB 2340375 and Alexa Fluor 594 donkey anti-rabbit 1:500, Jackson ImmunoResearch, AB 2340621) for 1 h at room temperature. Sections were then rinsed 3 more times for 10 min each in PBS and mounted on glass slides (SuperFrost Plus) and coverslipped after application of Fluoromount-G with DAPI (Southern Biotech).

Sections from the *Bmal1*-cKO^CeA^ experiment were stained in a separate run using the same cryoprotectant removal, blocking, wash, and mounting procedures. Primary antibodies were chicken-anti-GFP (1:10,000, Abcam ab13970) and rabbit-anti-BMAL1 (1:1,000, Novus Biologicals NB100-2288), applied overnight at 4 °C in blocking solution; secondary antibodies were Alexa Fluor 488 donkey anti-chicken (1:500, Jackson ImmunoResearch AB 2340375) and Alexa Fluor 594 donkey anti-rabbit (1:300, Invitrogen A21207), applied for 1 h at room temperature in PBS.

Images of slides consisting of the entire regions of interest were captured on a Keyence BZ-X710 fluorescence microscope at low magnification (2x). Higher magnification images (10x) of the center of the fiber tracts were captured to better evaluate their placement. Images were minimally processed to enhance brightness for representation purposes. For mice used in optogenetic experiments, we excluded three Tac1^PBN^, three Tac1^PBN→CeA^, and three Tac1^PBN→BNST^, mice that showed inadequate viral transduction or fiber tracts over the region of interest.

## Statistical analysis

Infrared (IR) activity counts were imported from ClockLab-style count files at 1-min resolution. Because experiments took place in constant dark conditions, the onset of each animals’ daily activity would occur at different times. As such, we anchored the daily start time of cyclic cues to activity onset on the last day of baseline recording, defined as circadian time 12 (CT12) throughout. In optogenetic experiments, CT12 was defined separately for each mouse based on its own activity onset, whereas in foot-shock experiments, CT12 was defined at the cohort level based on the group’s average activity onset phase. All subsequent analyses of IR activity and lever-press logs were performed using this CT-based time system rather than clock time.

### Actograms

Double-plotted actograms were generated from IR-derived foraging activity. For each animal, activity was plotted across consecutive 48-h windows by duplicating each day’s activity on the subsequent 24-h axis. Bars represented minute-by-minute foraging activity using the z-scored IR foraging activity. Maximum bar heights were set at 90% of maximum value to improve visual comparability across animals.

### Waveforms

Waveform plots were generated from IR-derived foraging activity using the 1-h rolling mean of z-scored counts, binned into 10-min intervals across the 24-h circadian cycle. For each animal, stage-specific analysis windows were defined as the last five days of the baseline stage, the last five days of the stimulation stage, and the first five days of the free-running phase. For each animal, activity was averaged within each 10-min bin and across each stage. Group-level 24-h waveforms were then plotted as the mean ± SEM across animals within each treatment group.

### Diurnality index

To quantify the temporal distribution of activity across the circadian cycle, we calculated a daily diurnality index from the IR-derived foraging signal using raw 1-min activity counts. For each animal, stage, and day, mean activity was computed separately for the subjective day (CT0–12) and subjective night (CT12–24). Daily percent diurnality was then calculated as:

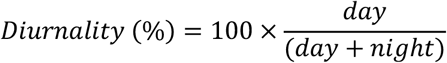

where day and night refer to the mean activity observed during the subjective day and subjective night, respectively. Under this formulation, 0% indicates absolute subjective nighttime activity and 100% indicates absolute subjective datytime activity. For each animal, stage-specific values were averaged across the same fixed stage-specific analysis windows described previously (baseline, stimulation, and free-running). For each treatment group we fit stage-specific models across the baseline, stimulation, and free-running windows and estimated within-group stage contrasts by estimated marginal means. To test for treatment-related shifts in diurnality, we then compared the diurnality from baseline to free-running stages of each treatment group against its control group (YFP controls for the optogenetic cohorts; wild-type mice for the *Bmal1*-cKO^CeA^ cohort) as an estimated-marginal-means contrast.

### Phase marker selection

To select an appropriate marker for evaluation of circadian phase, we first assessed four commonly used phase markers: center-of-gravity (COG), acrophase, activity onset, and M10 ^53,54^. Estimates of daily phases were plotted onto actograms and visually inspected for best fit. We selected COG as the principal phase marker for our analyses, because it produced a stable daily marker that could be readily visualized against the actogram structure and compared across experimental stages (Supp. Fig. 2).

Daily center-of-gravity (COG) phase estimates were computed from the IR-derived foraging signal using 1-min bins. For each animal, sensor, and day, daily COG was calculated by treating time of day as a circular variable over 24 h and weighting each time bin by its IR-derived activity. For each day, the daily 20th-percentile activity value was subtracted from the IR signal, and resulting negative values were set to zero before COG estimation. This yielded a daily COG phase estimate (expressed in h) and an associated mean resultant vector length (r), where larger r values indicate that activity was more tightly concentrated in time. Daily COG was used as the primary circadian phase marker for downstream circular analyses.

### Circular phase analysis and Rayleigh plots

Circular statistics were performed to test for significant clustering of daily activity phase markers. For each animal, daily COG times were collected from the baseline, stimulation, and free-running stages. For each animal within each stage window, a single-phase estimate was computed as the circular mean of the daily COG values. The within-animal mean resultant vector length (r) was also calculated as a descriptive measure of phase concentration across days. For each experimental group and stage window, we then calculated the circular mean direction, the mean resultant vector length, and the p value from the Rayleigh test of non-uniformity. The Rayleigh test was used to evaluate whether phase angles were significantly clustered rather than uniformly distributed around the 24-h cycle. To quantify uncertainty in the group mean direction, 95% confidence intervals were estimated by bootstrap resampling of animal-level phase values (2,000 resamples). In the Results, circular (Rayleigh) phase estimates are reported as the group circular mean CT ± the half-width of this bootstrap 95% confidence interval (mean ± 95% CI), together with the resultant length r; all linear summary statistics (phase-shift magnitude, diurnality, and bout counts) are reported as mean ± SEM.

To quantify persistent treatment-related shifts in circadian phase, baseline-normalized free-running phase shifts were calculated for each animal as the signed circular difference between the animal’s free-running phase estimate and its own baseline phase estimate. Group differences in these free-running – baseline phase shifts were then evaluated using two-sample permutation-based circular tests (package “Directional”^55^), comparing each treatment group against the control group (YFP controls for the optogenetic cohorts and wild-type mice for the *Bmal1*-cKO^CeA^ foot-shock cohort). P values were adjusted across treatment-versus-control comparisons using the Holm method.

### Lever-defined foraging episodes and bout structure

From the serial text logs outputted by the Arduino microcontroller (1-Hz resolution), we identified foraging bouts from timestamped microswitch press and release events together with the timing of optogenetic stimulation. For each animal and stage window, the total number of foraging bouts was calculated separately for the subjective day (CT0–12) and subjective night (CT12–24). Because control animals did not express ChR2, the maximum duration of each effective stimulation epoch in controls was limited to 6 s (compared to 2 min in opsin-expressing animals), which was chosen based on the average time required for opsin-expressing animals to retreat and terminate stimulation.

To quantify the probability that animals remained in the foraging area after stimulation onset, Kaplan– Meier curves were constructed from contiguous effective stimulation episodes, with time 0 defined as the onset of effective stimulation while the animal was actively foraging (microswitch pressed). The event was retreat from the foraging area, defined as termination of the microswitch-press state. Episodes in which animals did not retreat before the observed stimulation epoch ended were treated as right-censored. Survival curves were plotted separately by injection group using each group’s observed stimulation window.

To compare retreat probability across groups, Cox proportional hazards models were fit to stimulation episodes using control animals as the reference group. Because control animals had a shorter maximum stimulation duration than opsin-expressing animals, group comparisons were performed in a matched 6-s analysis window in which all episodes were right-censored at 6 s if retreat had not yet occurred. Hazard ratios therefore quantify the relative instantaneous hazard of retreat during the first 6 s after stimulation onset for each injection group relative to controls. Robust standard errors were clustered by animal to account for repeated episodes within mice.

## ACKNOWLEDGEMENTS

We thank Susan Phelps and Kit Mandeville for maintaining the mouse colony and lab members for their input during the development of this project.

## AUTHOR CONTRIBUTIONS

VYZ, SP, RDP, and HOD contributed to the concept. VYZ and SP performed and analyzed optogenetic experiments. KDD and VYZ performed and analyzed *Bmal1*-cKO^CeA^ experiments. JLP performed immunohistochemistry. VYZ wrote the initial manuscript with input from all authors.

## FUNDING

This work was supported by Howard Hughes Medical Institute and by the National Institutes of Health (grant number R01NS110012) and the Army Research Office (grant number W911NF-24-1-0013). The views and conclusions contained in this document are those of the authors and should not be interpreted as representing the official policies, either expressed or implied, of the Army Research Office or the U.S. Government. The U.S. Government is authorized to reproduce and distribute reprints for Government purposes, notwithstanding any copyright notation herein.

## CONFLICTS OF INTEREST

The authors declare no competing interests.

## SUPPLEMENTARY MATERIALS

**Figure 8.**
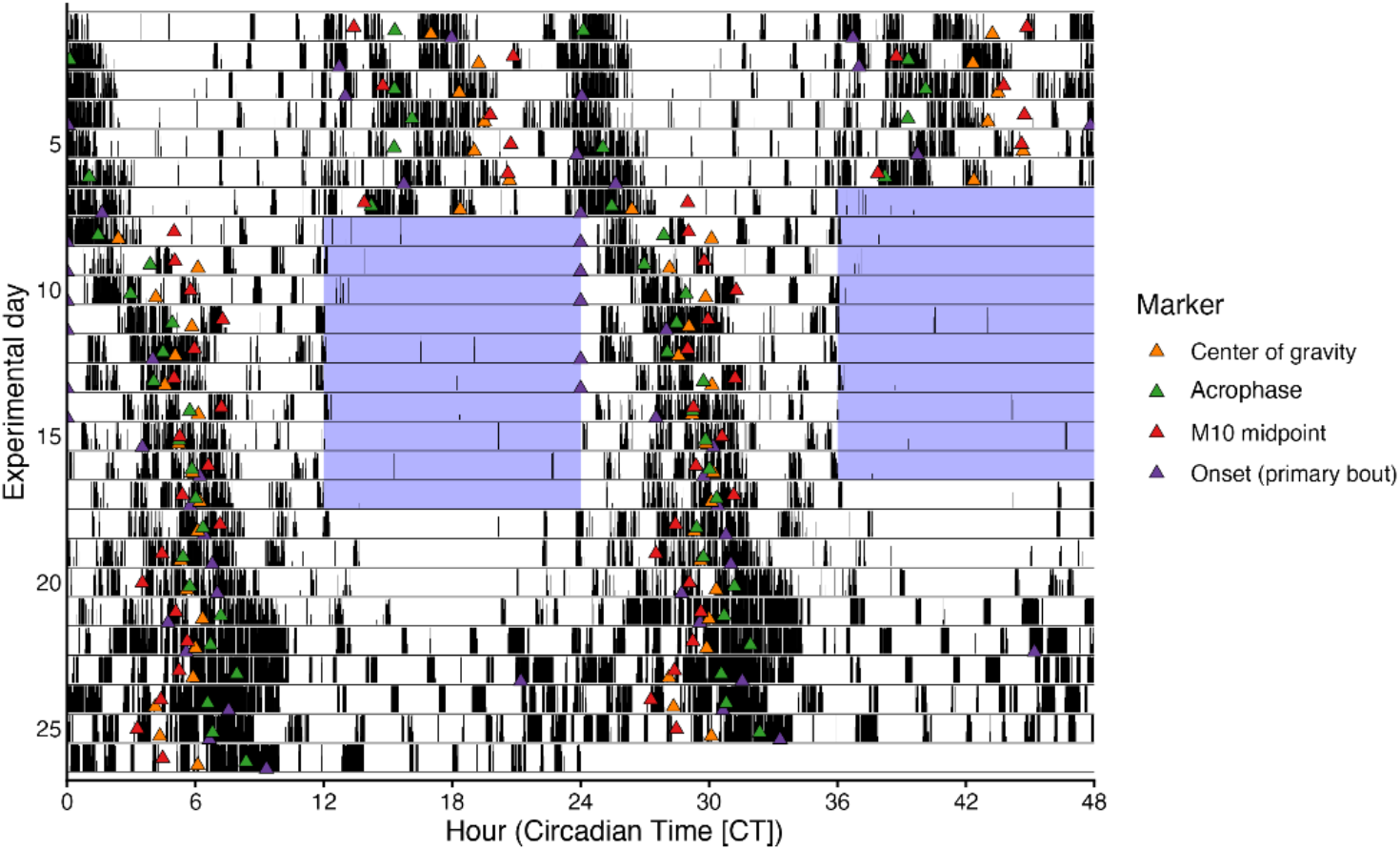
Foraging-bout frequency across optogenetic stimulation groups. Lever-defined foraging-bout frequency (bouts per day) during the subjective day (CT0–12) and subjective night (CT12–24) across baseline, stimulation, and free-run. Group mean ± SEM is shown overlayed with individual mice. n = 5 YFP control, 6 CGRP^PBN^, 6 Tac1^PBN^, 6 Tac1^PBN→CeA^ and 5 Tac1^PBN→BNST^ mice.

**Figure 8.**
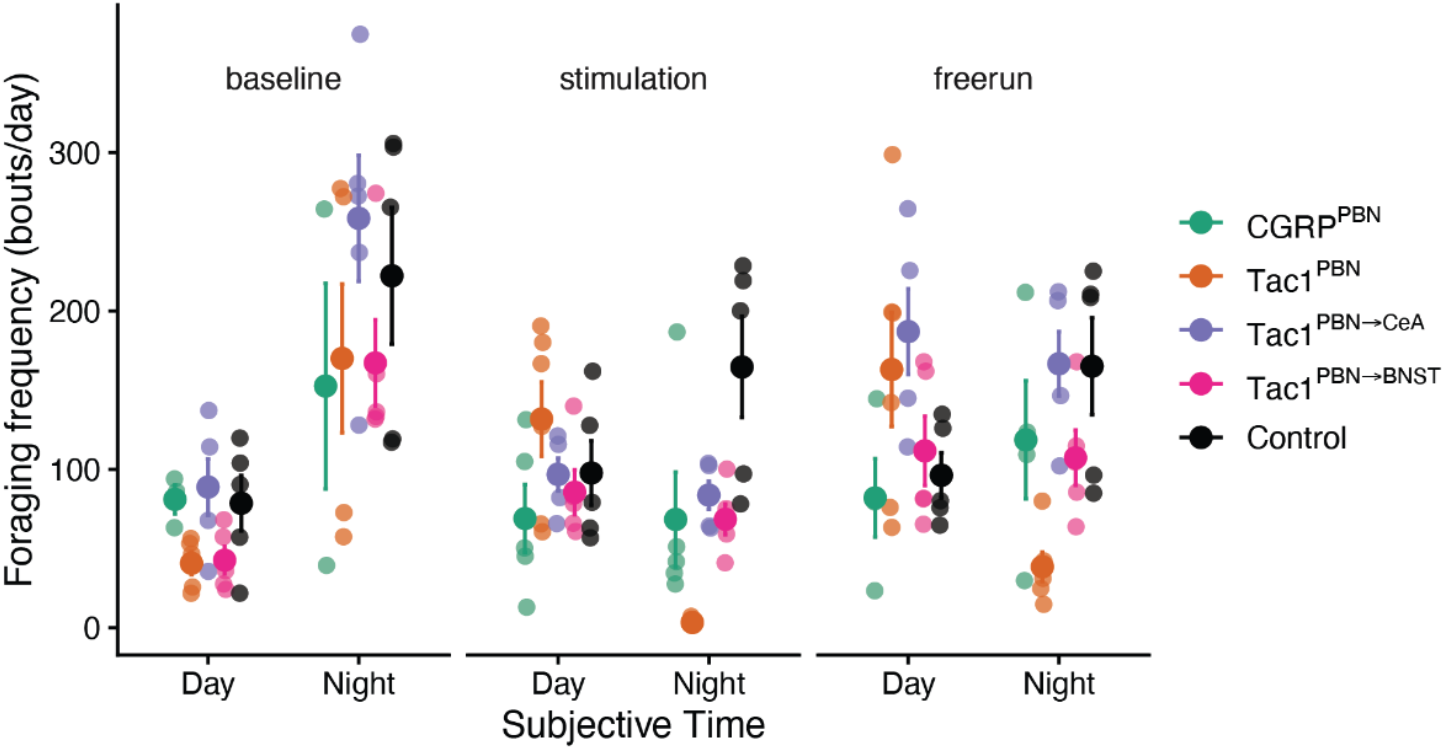
Comparison of candidate circadian phase markers. Representative double-plotted actogram of infrared foraging activity from a single Tac1^PBN^ mouse across baseline, optostimulation, and free-run, with four candidate daily phase markers overlaid: center of gravity (orange), acrophase (green), M10 midpoint (red) and onset of the primary activity bout (purple). Shaded regions mark the boundaries of the subjective-night (CT12–24) optostimulation window. Center of gravity produced a stable daily marker that could be readily visualized against the actogram structure and compared across stages and was selected as the principal phase marker for all subsequent circular analyses.

## Notes

### Competing Interest Statement

The authors have declared no competing interest.

